# Modeling confinement and reversibility of threshold-dependent gene drive systems in spatially-explicit *Aedes aegypti* populations

**DOI:** 10.1101/607267

**Authors:** Héctor M. Sánchez C., Jared B. Bennett, Sean L. Wu, Gordana Rašić, Omar S. Akbari, John M. Marshall

**Affiliations:** Division of Epidemiology and Biostatistics, School of Public Health, University of California, Berkeley, CA 94720, USA; Biophysics Graduate Group, University of California, Berkeley, CA 94720, USA; Mosquito Control Laboratory, QIMR Berghofer Medical Research Institute, Brisbane, Australia; Cell and Developmental Biology Section, Division of Biological Sciences, University of California, San Diego, CA 92093, USA; Innovative Genomics Institute, Berkeley, CA 94720, USA

## Abstract

**Background:** The discovery of CRISPR-based gene editing and its application to homing-based gene drive systems has been greeted with excitement, for its potential to control mosquito-borne diseases on a wide scale, and concern, for the invasiveness and potential irreversibility of a release. Gene drive systems that display threshold-dependent behavior could potentially be used during the trial phase of this technology, or when localized control is otherwise desired, as simple models predict them to spread into partially isolated populations in a confineable manner, and to be reversible through releases of wild-type organisms. Here, we model hypothetical releases of two recently-engineered threshold-dependent gene drive systems - reciprocal chromosomal translocations and a form of toxin-antidote-based underdominance known as UD^MEL^ - to explore their ability to be confined and remediated.

**Results:** We simulate releases of *Aedes aegypti*, the mosquito vector of dengue, Zika and other arboviruses, in Yorkeys Knob, a suburb of Cairns, Australia, where previous biological control interventions have been undertaken on this species. We monitor spread to the neighboring suburb of Trinity Park to assess confinement. Results suggest that translocations could be introduced on a suburban scale, and remediated through releases of non-disease-transmitting male mosquitoes with release sizes on the scale of what has been previously implemented. UD^MEL^ requires fewer releases to introduce, but more releases to remediate, including of females capable of disease transmission. Both systems are expected to be confineable to the release site; however, spillover of translocations into neighboring populations is less likely.

**Conclusions:** Our analysis supports the use of translocations as a threshold-dependent drive system capable of spreading disease-refractory genes into *Ae. aegypti* populations in a confineable and reversible manner. It also highlights increased release requirements when incorporating life history and population structure into models. As the technology nears implementation, further ecological work will be essential to enhance model predictions in preparation for field trials.

## Introduction

The discovery of CRISPR and its application as a gene editing tool has enabled gene drive systems to be engineered with much greater ease (1, 2). Recent attention has focused on homing-based drive systems and their potential to control mosquito-borne diseases on a wide scale, either by spreading disease-refractory genes (3) or by spreading genes that confer a fitness load or sex bias and thereby suppress mosquito populations (4, 5). The increased ease of gene editing has also advanced the entire field of gene drive, including systems appropriate during the trial phase of the technology (6). Such systems would ideally be capable of enacting local population control by: a) effectively spreading into populations to achieve the desired epidemiological effect, and b) being recallable from the environment in the event of unwanted consequences, public disfavor, or the end of a trial period.

As gene drive technology has progressed, a number of systems have been proposed with the potential to enact localized population control without spreading on a wide scale (6, 7). Sterile male releases provide one option (8), a recent version of which is based on the same molecular components as CRISPR gene drive systems (8, 9). At the interface between homing-based and localized suppression systems, an autosomal X chromosome-shredding system has been proposed that induces a transient male sex bias and hence population suppression before being selected out of the population (10). Population modification drive systems that display transient drive activity before being eliminated by virtue of a fitness cost, could also spread disease-refractory genes into populations in a localized way. Examples of this variety of drive system include split-gene drive (11), daisy drive (12) and killer-rescue systems (13). Each system has its own strengths and weaknesses, and could be suited to a different situation. In this paper, we theoretically explore the potential for two recently-engineered threshold-dependent gene drive systems to achieve localized and reversible population modification in structured populations - reciprocal chromosomal translocations (14) and a toxin-antidote-based system known as UD^MEL^ (15).

Threshold-dependent gene drive systems must exceed a critical threshold frequency in a population in order to spread. Based on this dynamic, simple population models, in which two randomly mating populations exchange small numbers of migrants with each other, predict that these systems can be released at high frequencies in one population and spread to near-fixation there, but never take off in the neighboring population because they do not exceed the required threshold there (16, 17). These systems can also be eliminated through dilution to sub-threshold levels with wild-type organisms at the release site, making them excellent candidates for the trial phase of a population modification gene drive strategy, or when localized population modification is desired (16). Elimination of a non-driving transgene can in fact be more difficult, as the dynamics of threshold-dependent systems actively drive them out of populations at sub-threshold levels. However, whether these dynamics hold in real ecosystems depends crucially on the dispersal patterns and population structure of the species being considered. First steps towards modeling the spatial dynamics of these systems have been taken by Champer *et al.* (18), who model spatially-structured releases of various threshold-dependent systems without life history, and Huang *et al.* (19), who model engineered underdominance (20) on a grid-based landscape incorporating life history for *Aedes aegypti*, the mosquito vector of dengue, Zika and other arboviruses.

Here, we present a detailed ecological analysis of the expected population dynamics of two recently-engineered threshold-dependent drive systems, translocations and UD^MEL^, in *Ae. aegypt* in a well-characterized landscape – Yorkeys Knob, a suburb ~17 km northwest of Cairns, Australia (Figure 1C) – suitable for confineable and reversible releases. Yorkeys Knob and the nearby town of Gordonvale were field sites for releases of *Wolbachia*-infected mosquitoes in 2011 (21) and the prevalence of *Wolbachia* infection over time provided information on the number of adult *Ae. aegypti* mosquitoes per household and other mosquito demographic parameters for that location (22), as well as an opportunity to validate our modeling framework. *Wolbachia* is an intracellular bacterium and biocontrol agent that biases inheritance in its favor if infected females are present, and blocks transmission of dengue and other arboviruses (23). Yorkeys Knob is a partially isolated suburb, separated by a 1-2 km-wide uninhabited, vegetated area from the nearest suburb, Trinity Park, a control site for the *Wolbachia* trial. This allowed us to simulate trials of transgenic mosquitoes in a well-characterized population, while also theoretically exploring their potential of spread to a neighboring community.

**Figure 1.**
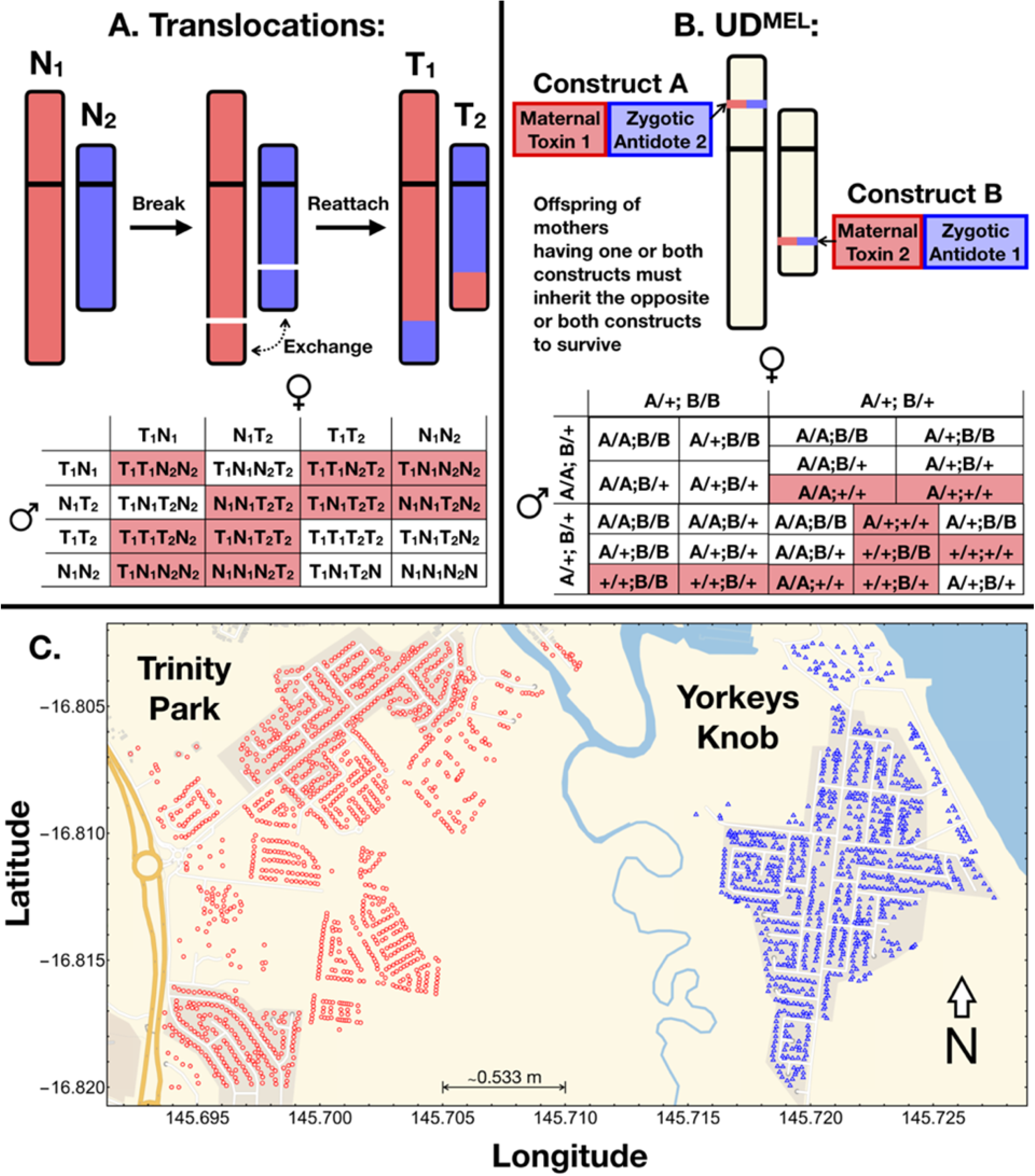
Inheritance and landscape features of the modeling framework. *(A)* Reciprocal translocations (T1 and T2) result from the mutual exchange between terminal segments of two non-homologous chromosomes (N1 and N2). The cross here depicts possible parental gametes, with respect to the translocation, and offspring that result from matings between them. Matings between wild-type organisms or translocation homozygotes result in viable offspring; but translocation heterozygotes produce unbalanced gametes, and many of the resulting offspring are unviable (shaded). This results in a heterozygote disadvantage and threshold-dependent population dynamics. *(B)* UD^MEL^ is composed of two unlinked constructs (here referred to as A and B), each consisting of a maternally-expressed toxin and a zygotically-expressed antidote for the toxin on the opposite construct. The cross here represents matings between two of the nine possible parental genotypes (“+” represents the wild-type allele, and “A” and “B” represent alleles corresponding to the two UD^MEL^ constructs). The complete inheritance pattern is depicted in Figure S1. Offspring lacking the antidotes to the maternal toxins produced by their mother are unviable (shaded). At high population frequencies, the selective advantage on the constructs, by virtue of the antidotes, outweighs the fitness load due to the toxins, and hence results in frequency-dependent spread. *(C)* Distribution of households in Yorkeys Knob (blue) and Trinity Park (red) in Cairns, Queensland, Australia. Households serve as individual *Aedes aegypti* populations in our metapopulation framework, with movement of adult *Ae. aegypti* between them. Yorkeys Knob serves as a simulated release site, and Trinity Park as a simulated control site.

A number of other ecological details are relevant to the spread of threshold-dependent gene drive systems that have not been considered in previous modeling studies. Perhaps of greatest importance, the frequency of the introduced transgene in the mating pool is markedly different from the frequency of introduced adults. It is typical to release only male mosquitoes as part of an intervention, as only females are involved in human disease transmission. Life cycle and mating structure therefore become relevant, as immature life stages are not available for mating, and female adults are thought to mostly mate only once soon after emergence (24). This means that many of the released adult males will not find a mating partner, and hence larger releases will be required to exceed threshold frequencies than predicted in simple population frequency models.

The nature of mosquito dispersal behavior is also relevant to the spatial dispersal of transgenes. Our species of interest, *Ae. aegypti*, is understood to display leptokurtic dispersal behavior in a suburban setting, in which mosquitoes tend to remain in the same household for the majority of their lifespan, while a few mosquitoes disperse over larger distances (25). With these landscape, dispersal and life cycle considerations accounted for, we theoretically explore the ability to drive two threshold-dependent systems, translocations and UD^MEL^, into populations of *Ae. aegypti* in one community, Yorkeys Knob, without them spreading in significant numbers to a neighboring community, Trinity Park, and to be remediated from Yorkeys Knob at the end of the simulated trial period.

## Results

### Model framework

We use the Mosquito Gene Drive Explorer (MGDrivE) modeling framework (26) to model the spread of translocations and UD^MEL^ through spatially-structured mosquito populations (Figure 1). This is a genetic and spatial extension of the lumped age-class model of mosquito ecology (27) modified and applied by Deredec *et al.* (28) and Marshall *et al.* (29) to the spread of homing gene drive systems. The framework incorporates the egg, larval, pupal and adult life stages, with egg genotypes being determined by maternal and paternal genotypes and the allelic inheritance pattern of the gene drive system. Spatial dynamics are accommodated through a metapopulation structure in which lumped age-class models run in parallel and migrants are exchanged between populations according to a zero-inflated exponential dispersal kernel with parameters defined in Table S1. Further details of the framework are described in the Materials and Methods section.

Applying the MGDrivE modeling framework to our research questions, we incorporate the inheritance patterns of reciprocal chromosomal translocations and UD^MEL^ into the inheritance module of the model (Figure 1A-B, Figure S1), the life cycle parameters of *Aedes aegypti* (Table S1) into the life history module, and the distribution of households in Yorkeys Knob (923 households) and Trinity Park (1301 households) along with their expected mosquito population sizes and movement rates between them into the landscape module (Figure 1C). The suburb of Trinity Park served as a control site for field releases of *Wolbachia*-infected mosquitoes, to quantify the extent to which the *Wolbachia* infection could spread from one community to another, and plays a similar role for our simulated releases of threshold-dependent gene drive systems.

The inheritance patterns that result from chromosomal translocations are depicted in Figure 1A. Chromosomal translocations result from the mutual exchange between terminal segments of two nonhomologous chromosomes. When translocation heterozygotes mate, several crosses result in unbalanced genotypes and hence unviable offspring, resulting in a heterozygote reproductive disadvantage. This results in bistable, threshold-dependent population dynamics, confirmed in laboratory drive experiments (14). The inheritance patterns produced by the UD^MEL^ system are depicted in Figure 1B. This system consists of two unlinked constructs, each possessing a maternally-expressed toxin active during oogenesis, and a zygotically-active antidote expressed by the opposite construct. The resulting dynamic, confirmed in laboratory drive experiments (15), is gene drive for allele frequencies greater than ~24%, in the absence of a fitness cost.

### Model validation

Using data from field trials of *Wolbachia*-infected *Ae. aegypti* mosquitoes in Yorkeys Knob and Gordonvale, Australia (21), we validated our modeling framework prior to application to other threshold-dependent drive systems. *Wolbachia* biases the offspring ratio in favor of those carrying *Wolbachia* through a mechanism known as cytoplasmic incompatibility, in which offspring of matings between infected males and uninfected females result in the death of some or all progeny, while matings involving infected females tend to produce infected offspring (30). For the *wMel* strain of *Wolbachia* that was used in the Australian field trials, incompatible crosses produce no viable offspring, and *Wolbachia* is inherited by all offspring of infected females. There is also a fitness cost associated with *Wolbachia* infection, the value of which has been estimated between 0-20% (21, 31, 32).

Data in Figure 1 of Hoffmann *et al.* (21) provides information on the size and timing of releases of *Wolbachia*-infected mosquitoes, as well as the population frequency of *Wolbachia* infection over time. Release data informed the simulation parameters, and surveillance data was used to compare to model predictions for a range of biological parameter values, including baseline adult mortality rate and *Wolbachia*-associated fitness cost. Weekly releases of 20 *Wolbachia*-infected mosquitoes (10 female and 10 male) per household at a coverage of 30% were modeled over 10 weeks in the spatially-explicit landscapes of Yorkeys Knob and Gordonvale with the exception that, in Gordonvale, the fifth release was postponed by a week due to a tropical cyclone (21). Based on the comparative plots of *Wolbachia* surveillance data alongside model predictions in Figure S2, we found that model predictions most closely matched field data for a baseline adult mortality rate of 0.090 per day (33), as compared to a mortality rate of 0.050 per day (34), and that while observations matched predictions quite well for both 5% and 10% fitness costs, a 10% fitness cost is closer to that estimated elsewhere (21, 31, 32). Model predictions in Figure 2 use these parameter values, along with others listed in Table S1, and provide a good validation of our modeling framework.

**Figure 2.**
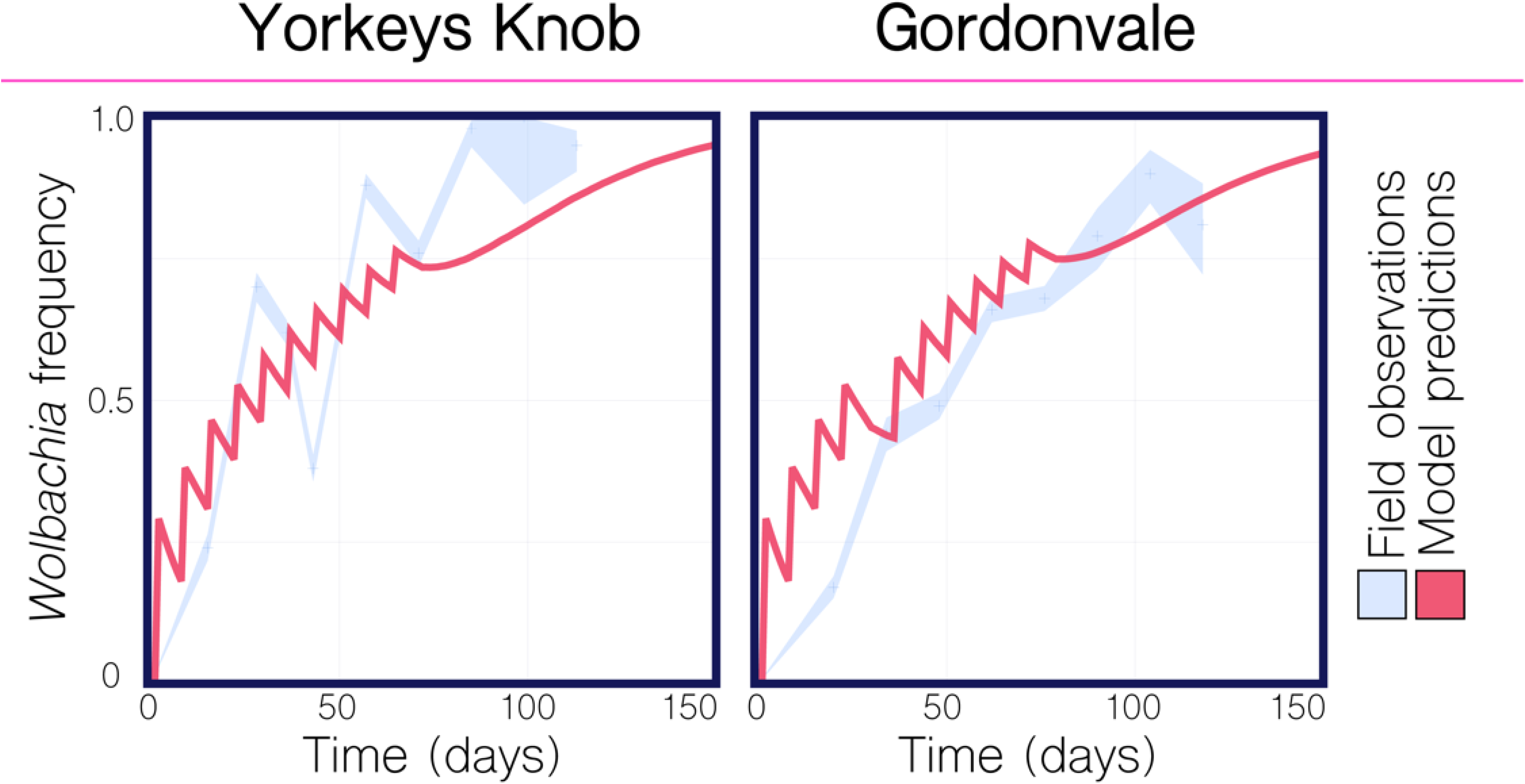
Comparison of model predictions to *Wolbachia* field trial data from Yorkeys Knob and Gordonvale, Australia. Field observations of *Wolbachia* population frequency are depicted in light blue, with 95% binomial confidence intervals based on the frequency and sample size reported for the 2011 field trials in Yorkeys Knob and Gordonvale (21). Model predictions are depicted for an analogous release scheme consisting of 20 *Wolbachia*-infected mosquitoes (10 female and 10 male) per household at a coverage of 30% over 10 weeks with the exception that, in Gordonvale, the fifth release was postponed by a week due to a tropical cyclone. Parameter values listed in Table S1. *Wolbachia* infection is associated with a 10% fitness cost. Agreement between observations and predictions is strong, providing good validation of the modeling framework.

### Population replacement and remediation for translocations

The use of translocations for transforming pest populations was initially suggested by Serebrovskii (1940) (35) and later Curtis (1968) (36) for the introduction of disease-refractory genes into mosquito populations. A number of models have been proposed to describe their spread through randomly-mating populations (14, 16, 37, 38); however, with one recent exception addressing spatial structure (18), these have largely ignored insect life history and mating structure. Such models suggest that the translocation need only exceed a population frequency of 50%, in the absence of a fitness cost associated with the translocation, to spread to fixation in a population, which could conceivably be achieved through a single seeding release round. Here, we find that incorporating life history and population structure into mosquito population dynamic models significantly increases release requirements.

In Figures 3–4, based on the precedent set by the 2011 *Wolbcahia* field trial (21), we consider weekly releases of 20 adult *Ae. aegypti* males homozygous for the translocation for given durations and coverage levels, where coverage level is the proportion of households that receive the releases. Releases are simulated in the community of Yorkeys Knob, in which prior releases of *Wolbachia*-infected mosquitoes suggested a local population of ~15 adult *Ae. aegypti* per household (22), and for mosquito movement rates inferred from previous studies (25, 39, 40) (Table S1). For a coverage level of 100%, and in the absence of a fitness cost, four weekly releases of 20 *Ae. aegypti* males (~3:1 release to local males) are required for the translocation to spread to fixation in the community (Figure 3), as opposed to the single release expected when ignoring life history and population structure (37). As coverage is reduced to 50%, the required number of releases increases to 7, and for a coverage level of 25%, as seen for the World Mosquito Program in Yorkeys Knob, the required number of releases increases to 16 (Figures 3–4). Although large, these releases are achievable, considering the much larger releases conducted for sterile insect programs (41).

**Figure 3.**
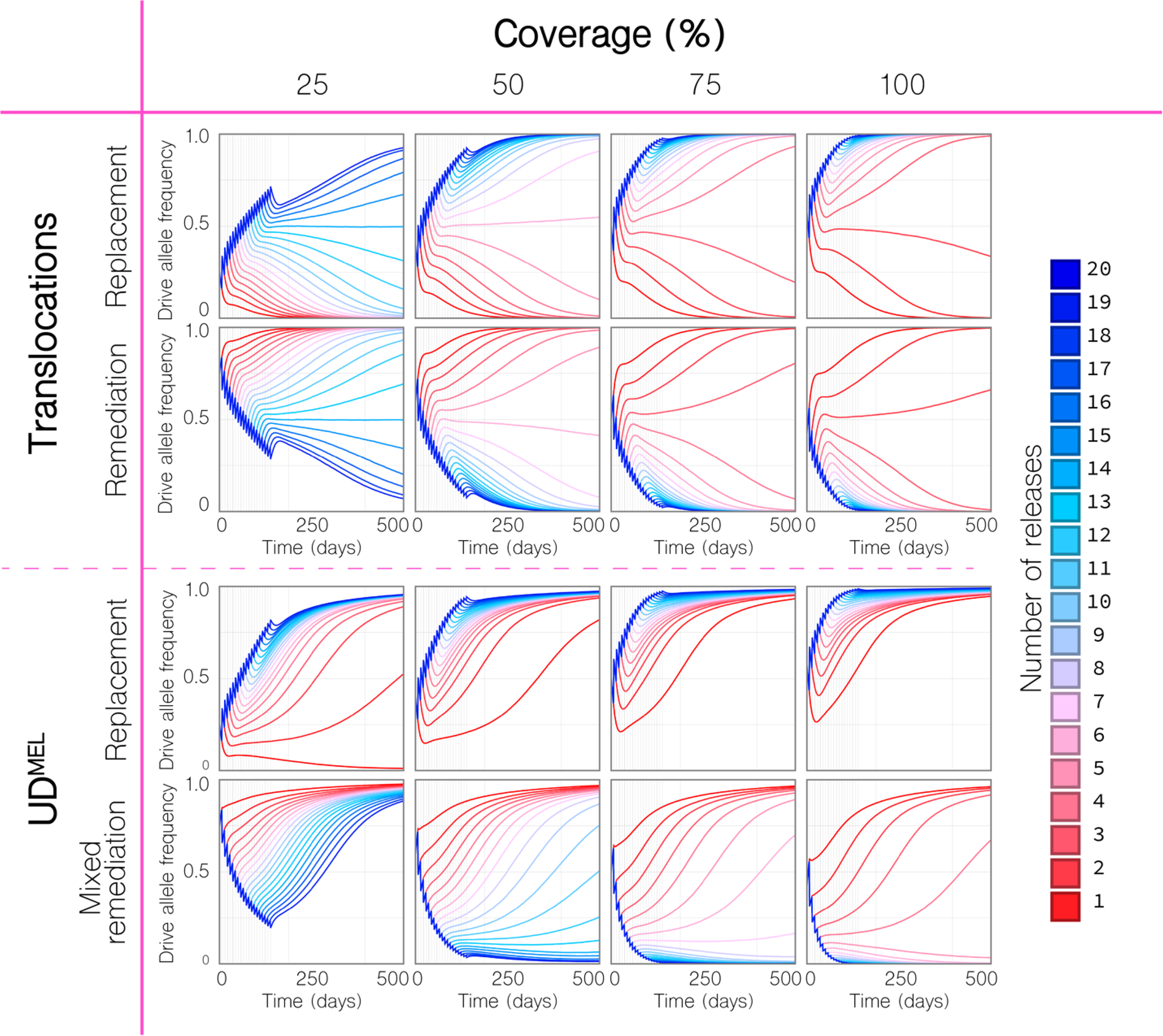
Replacement and remediation results for translocations and UD^MEL^. Time-series results are shown for a given number of weekly releases of 20 adult *Ae. aegypti* per household with the intent of population replacement or remediation in the community of Yorkeys Knob (Figure 1C), and at given coverage levels, where coverage is the proportion of households that receive the releases. For population replacement, releases are of males homozygous for the translocation or UD^MEL^ into a wild-type population. For remediation of translocations, releases are of wild-type males into a population homozygous for the translocation. For UD^MEL^, remediation is not possible through releases of males only, and so “mixed remediation” is considered, in which releases consist of 10 females and 10 males. *(Top)* Replacement and remediation are symmetric for translocations. At a coverage of 50%, seven or more releases result in the translocation being driven to fixation or remediated from the population. *(Bottom)* Release requirements for UD^MEL^ are smaller for population replacement, but larger for mixed remediation. At a coverage of 50%, a single release results in UD^MEL^ being driven to fixation.

To simulate remediation of a translocation, we consider weekly releases of 20 adult *Ae. aegypti* wild-type males in the community of Yorkeys Knob, whereby the translocation has already reached fixation in that community. In the absence of a fitness cost associated with the translocation, translocations are symmetrical in their threshold dynamics and so, for a coverage level of 100%, four weekly releases are required for the translocation to be completely remediated from the community, and for a coverage of 25%, 16 weekly releases are required for the translocation to be completely remediated (Figures 3–4). Encouraging features of these results are that: i) remediation can be achieved through releases of non-biting, non-disease-transmitting males, ii) release sizes are achievable, and iii) despite the spatial household structure, both replacement and remediation are complete within the community. The time to replacement is highly dependent on the coverage level and number of releases; but is reasonably quick given sufficient releases. At a coverage of 50%, 20 weekly releases led to the translocation spreading to a frequency >99% within half a year of the final release (or within 300 days of the first release). For equivalent wild-type releases, this is the same as the time to >99% elimination.

### Population replacement and remediation for UD^MEL^

UD^MEL^ was the first synthetic gene drive system to be engineered that displays threshold-dependent dynamics (15). The system consists of two unlinked constructs, each possessing a maternally expressed toxin active during oogenesis, and a zygotically active antidote expressed by the opposite construct (Figure 1B). At low population frequencies, the maternal toxin confers a significant selective disadvantage, leading to elimination, while at high population frequencies, the zygotic antidote confers a selective benefit in the context of a prevalent toxin, leading to fixation. The dynamics of this system in randomly-mating populations have been characterized by Akbari *et al.* (15), suggesting that the system need only exceed a population frequency of ~24%, in the absence of a fitness cost, to spread to fixation, while the wild-type must exceed a population frequency of ~76% to eliminate the construct. Both replacement and remediation should therefore be achievable with 1-2 releases of transgenic and wild-type organisms, respectively; however, as for translocations, we find that incorporating life history and population structure into our models increases release requirements in both cases.

In Figures 3–4, we consider weekly releases of 20 adult *Ae. aegypti* males homozygous at both loci for the UD^MEL^ system in the community of Yorkeys Knob. The lower threshold for UD^MEL^ as compared to translocations means that replacement is much easier to achieve for UD^MEL^. For a coverage level of 50% or higher, and in the absence of a fitness cost, a single release of 20 *Ae. aegypti* males leads to the UD^MEL^ system spreading to fixation throughout the community (Figure 3). As coverage is reduced to 25%, the required number of releases to achieve fixation increases to two (Figures 3–4). As for translocations, the time to replacement is highly dependent on the coverage level and number of releases. From Figure 3, it is apparent that UD^MEL^ reaches total allele fixation slowly, although the number of individuals having at least one copy of the transgene increases quickly. At a coverage of 50%, 10 weekly releases lead to wild-type individuals falling to a frequency <2% within 2.6 years of the final release (or within 3 years of the first release).

Remediation, however, is more difficult to achieve for UD^MEL^ compared to translocations due to the higher threshold that wild-type organisms must exceed to eliminate UD^MEL^. Additionally, wild-type females must be included in the releases to propagate the wild-type allele because, assuming continued functioning of UD^MEL^ components, the maternal toxins of females having UD^MEL^ at both loci kill all offspring that do not inherit UD^MEL^ at both loci. To simulate remediation, we first consider weekly releases of 10 adult *Ae. aegypti* wild-type females and 10 adult males in the community of Yorkeys Knob. In the absence of a fitness cost associated with the UD^MEL^ construct, and for a coverage level of 75%, nine weekly releases are required for a reduction in UD^MEL^ allele frequency over the first year (Figure 3); however a closer inspection of the simulation results reveals that complete remediation of UD^MEL^ from the community is not possible even with 20 releases, as the UD^MEL^ allele frequency bounces back. Comparison of these results to those for a panmictic population with a population size equal to that of Yorkeys Knob reveals that complete remediation of UD^MEL^ can occur at coverage levels as low as 25% (for 16 or more weekly releases) (Figure S3). Inspection of the spatially-explicit simulation results suggests that the rebound in UD^MEL^ allele frequency in the structured population is due to UD^MEL^ remaining at super-threshold levels after the wild-type releases in a small number of households, and slowly recolonizing the landscape following that. Complete remediation of UD^MEL^ is possible, however, for 15 or more releases at a coverage level of 100% (Figure 4). These results make a strong case for translocations as preferred systems to introduce transgenes in a local and reversible way as: i) remediation of UD^MEL^ requires releases of biting, vector-competent females, and ii) release requirements for these biting, vector-competent females are burdensomely high due to the hig threshold that must be surpassed consistently throughout a spatially structured population.

**Figure 4.**
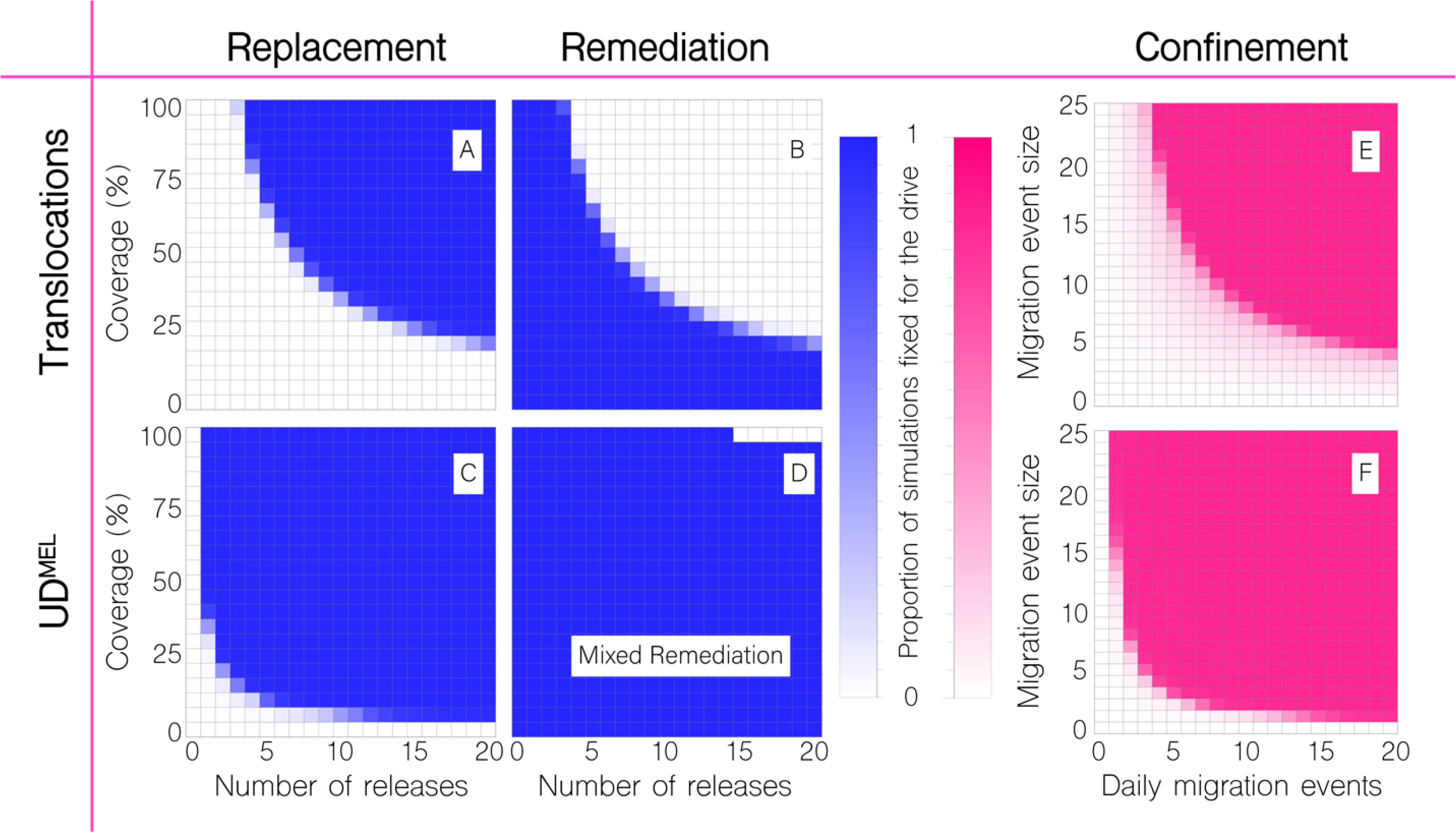
Replacement, remediation and confinement outcomes for translocations and UD^MEL^. Outcomes are depicted for the proportion of 50 stochastic simulations of population replacement, remediation and confinement of translocations and UD^MEL^ that result in fixation of each system. *(A-D)* For replacement and remediation, each cell corresponds to a given number of releases (horizontal axis) and coverage level (vertical axis), given 20 adult *Ae. aegypti* per household per release. For replacement, releases are of males homozygous for the system into a wild-type population. For remediation of translocations, releases are of wild-type males into a population homozygous for the translocation, and for mixed remediation of UD^MEL^, releases are of wild-type females and males into a population homozygous for UD^MEL^. Blue cells represent cases where all simulations result in fixation of the system, and white cells represent cases where the wild-type is fixed in all simulations. *(G-H)* For confinement, each cell corresponds to a daily number of batch migration events (horizontal axis) of a given size (vertical axis) from Yorkeys Knob, where the system is fixed, to Trinity Park, where the system is initially absent. White cells represent cases where all simulations result in fixation of the system in Trinity Park, and dark pink cells represent cases where the wild-type is fixed in all simulations. These results are encouraging for translocations as systems for introducing transgenes in a local and reversible way as: i) they can be remediated through an achievable number of male-only releases, and ii) they require more batch migration events to spread to neighboring communities.

### Confinement of translocations & UD^MEL^ to release site

Confinement of translocations and UD^MEL^ to partially-isolated populations has previously been modeled by Marshall & Hay (16) and Akbari *et al.* (15). In both cases, two randomly-mating populations were modeled that exchange migrants at given rates. Population structure was otherwise ignored, as was mosquito life history. Results from these analyses suggest that translocations would spread and remain confined to populations for which the migration rate is less than ~5.8% per mosquito per generation (16), and that UD^MEL^ would remain confined to populations for which the migration rate is less than ~1.6% per mosquito per generation (15). These migration rates are relatively low, however this may be beneficial for the types of landscapes we are considering here, whereby the system may spread between neighboring households, but not from one suburb to another. Recently, Champer *et al.* (18) showed that translocations would remain confined to and persist in a population connected to another by a “migration corridor” under a range of parameter values.

For our landscape of interest - the suburbs of Yorkeys Knob and Trinity Park - it is very unlikely that *Ae. aegypti* mosquitoes will travel from one suburb to another by their own flight. Extrapolating the exponential dispersal kernel used in our simulations, fitted to data from mark-release-recapture experiments collated by Guerra *et al.* (39), suggests these events to be negligible, before accounting for the fact that the intervening vegetated area may serve as a barrier to *Ae. aegypti* flight (42). Furthermore, rare migrant mosquitoes are unlikely to cause the threshold frequency for either drive system to be exceeded, thus making spatial spread due to such movements unlikely. In considering confineability to the release suburb, we therefore model “batch migration,” in which several mosquitoes are carried, perhaps by a vehicle, from one community to another at once. Batch migration events could be thought of as several adult mosquitoes being carried at once, or perhaps more likely, as a larval breeding site, such as a tyre, being carried from one household to another, with several adults emerging from the tyre following transport. We model batch migration events as occurring between randomly chosen households, and vary the number of daily migration events and the effective number of adults carried per event. For computational simplicity, we focus on migration events from Yorkeys Knob, in which either system has already reached fixation, to households in Trinity Park, which is initially fixed for wild-type mosquitoes.

In Figure 4E-F, we see that both the number and size of daily batch migration events affect the chance of either system establishing itself in the neighboring suburb, Trinity Park. For translocations, ~16 daily migration events of batches of 5 adults are required for spread in Trinity Park. For batches of 10 adults, ~9 daily migration events are required, and for batches of 20 adults, ~5 daily migration events are required. For UD^MEL^, ~3 daily migration events of batches of 5 adults are required for spread in Trinity Park, and for batches of 10 adults, ~2 daily migration events are required.

These results continue to make a strong case for translocations as preferred systems to introduce transgenes in a local and reversible way as: i) many more batch migration events are required to lead to spread for translocations as opposed to UD^MEL^, and ii) the rate of migration events required for translocations to spread is higher than what would be expected between these communities. Specifically, *Wolbachia* releases in Yorkeys Knob in 2011 provide evidence for occasional batch migrations to the nearby suburb of Holloways Beach; however the spatio-temporal pattern of *Wolbachia* spread, as inferred from monitored trap data, suggests only ~1-2 batch migration events over the course of a month, consisting of less than 5 adult females per event (21).

### Sensitivity analysis

A theoretical study by Khamis *et al.* (43) on toxin-antidote-based underdominance gene drive systems, similar to UD^MEL^ but for which the toxins are zygotic rather than maternal (20), found that the gene drive threshold frequency is highly sensitive to: i) the increase in adult mortality rate in organisms having the transgene, ii) the duration of the larval life stage, and iii) parameters determining the character or strength of larval density dependence. In Figure 5 and Figure S4, we explore the sensitivity of our model outcomes of replacement, remediation and confinement for translocations and UD^MEL^ as we vary: i) the duration of the larval life stage, ii) the baseline adult mortality rate, iii) the fitness cost associated with the gene drive system, and iv) the mean adult dispersal distance. For translocations, we model a 10% fitness cost as a 10% reduction in mean adult lifespan for organisms homozygous for the translocation, and a 5% reduction for organisms heterozygous for the translocation. For UD^MEL^, since its inheritance bias is induced through the action of maternal toxins, we model a 10% fitness cost as a 10% reduction in female fecundity for organisms homozygous for UD^MEL^ at both loci, with 2.5% additive fitness costs contributed by each transgenic allele.

**Figure 5.**
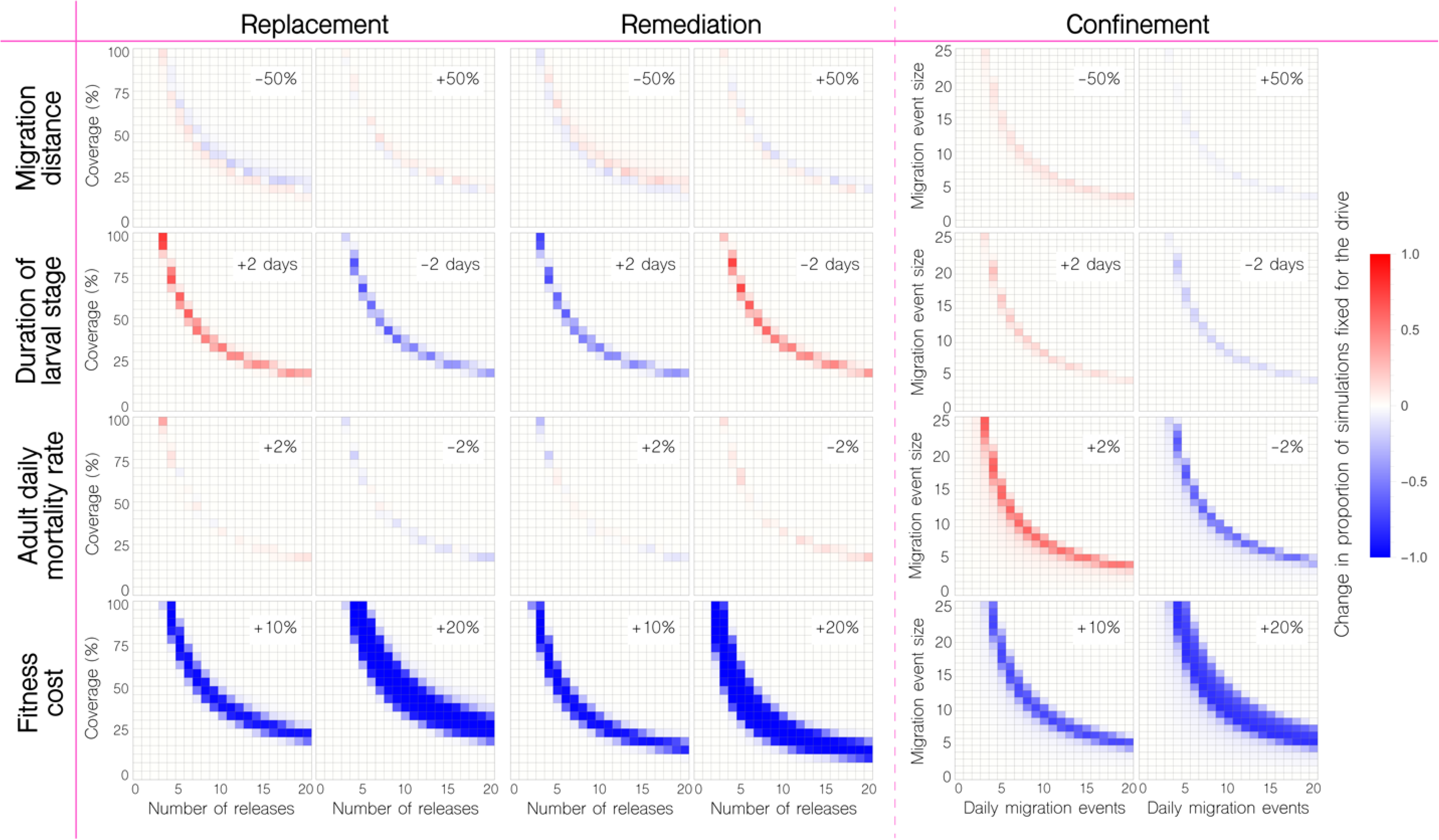
Sensitivity of model outcomes for replacement, remediation and confinement of translocations. Changes are depicted in the proportion of 50 stochastic simulations that result in fixation for replacement, remediation and confinement of translocations. Proportions are compared to those in the first row of Figure 4 as we vary: i) the mean dispersal distance of adult mosquitoes (+/− 50%), ii) the duration of the larval life stage (+/− 2 days), iii) the baseline adult mortality rate (+/− 2%), and iv) the fitness cost associated with being homozygous for the translocation (+10% or +20%). Fitness costs have the greatest impact on the release scheme required for the system to be fixed or remediated from the population, given the life parameters considered. Fitness costs also lead to more batch migration events being required for invasion of Trinity Park. A small increase in the baseline adult mortality rate leads to slightly fewer batch migration events being required for invasion of Trinity Park; however, comparison to migration rates inferred from field data suggests that confinement is still expected.

For translocations, the associated fitness cost had the greatest impact on the release scheme required for the system to be fixed or remediated from the population, given the life parameters considered (Figure 5). A 10% fitness cost led to ~10 weekly releases at a coverage of 50% being required for the translocation to reach fixation (an increase of 3 releases), while a 20% fitness cost led to ~13 weekly releases being required (an increase of 6 releases). Small changes in the duration of the larval life stage had minor impacts on the release requirements, with an increase in larval lifespan of 2 days leading to one more weekly release being required for the translocation to reach fixation, and vice versa. A 2% change in the baseline adult mortality rate and 50% change in the mean migration distance had negligible impact on release requirements. Remediation, on the other hand, requires fewer wild-type releases when there is a fitness cost associated with the translocation. A 10% fitness cost led to 5 weekly releases at a coverage of 50% being sufficient to eliminate the translocation (a decrease of 2 releases), and a 20% fitness cost led to 4 weekly releases at a coverage of 50% being sufficient for elimination (a decrease of 3 releases). Small changes in the duration of the larval life stage had minor impacts on the wild-type release requirements for elimination, with an increase in larval lifespan of 2 days or a 2% decrease in the adult mortality rate leading to one fewer release being required.

The sensitivity of our predictions regarding confinement to the release site are of particular interest, as invasion of a neighboring community may be more likely under some parameter values than others. Fortunately, a fitness cost associated with the translocation leads to a higher threshold, and hence more batch migration events required for invasion of a neighboring community. A 10% fitness cost led to ~2-3 additional daily migration events of 10 adults required for spread to Trinity Park, and a 20% fitness cost led to ~6-7 additional daily migration events required (Figure 5). Also noteworthy, a 2% increase in the adult mortality rate led to ~2 fewer daily migration events required for spread to Trinity Park - i.e. ~7 migration events for batches of 10 adults, and ~14 migration events for batches of 5 adults. While still above inferred batch migration rates, this highlights that there could exist parameter sets beyond those explored for which invasion is feasible.

UD^MEL^ displays similar parameter sensitivities regarding fixation, remediation and batch migration outcomes as for translocations, with the exception that these outcomes are less sensitive to fitness costs (Figure S4), likely due to the fact that fitness is accommodated through a reduction in female fecundity rather than an increase in adult mortality. A 20% fitness cost led to ~1 additional weekly release being required for the system to spread to fixation, whether at a coverage of 25% or 50%. Similarly, for an invasion of Trinity Park, a 10% fitness cost required ~1 additional daily migration event of 5 adults, and a 20% fitness cost required ~2 additional daily migration events. Of note, a 2% increase in the adult mortality rate or a 2 day increase in the duration of the larval stage led to ~1 fewer daily migration event required for spread to Trinity Park, making this now very achievable - i.e. ~2-3 migration events for batches of 5 adults, and ~1-2 migration events for batches of 10 adults.

Finally, we conducted an analysis of the sensitivity of our results to population structure, exploring the impact of: i) removing all population structure by treating Yorkeys Knob and Trinity Park as randomly mixing populations and, ii) incorporating heterogeneity in mosquito household population size. Results of the comparison to panmictic populations are depicted in Figure S5. Previously, we had seen that introducing population structure greatly increases the release requirements to eliminate UD^MEL^ from a community (Figure S3). The trend of higher release requirements in structured populations is also seen for translocations, although to a lesser extent, with one additional release required for either replacement or remediation at a coverage of 75%, two additional releases required at a coverage of 50%, and 7-8 additional releases required at a coverage of 25%. Invasion of a neighboring population, on the other hand, requires ~1-2 fewer daily migration events in structured populations for batches of 10 adults having either the translocation or UD^MEL^.

Incorporating heterogeneity in household mosquito population size, we retain a mean of 15 adults, as inferred from *Wolbachia* field trial data in Yorkeys Knob (22), and distribute population sizes across households according to a zero-inflated, truncated exponential distribution with 55% of households having no mosquitoes, and none having more than 45 adults. This distributional form, including zero inflation, is based on results of a large field survey conducted across a set of households in Kamphaeng Phet province, Thailand (44).

Including this source of heterogeneity substantially increases release requirements for replacement and remediation with translocations, with 3-4 additional releases required at a coverage of 75%, 5-6 additional releases required at a coverage of 50%, and 20 releases being insufficient at a coverage of 25% (Figure S6). Release requirements are marginally increased for UD^MEL^, with ~1-2 additional releases required at coverages of 25-100%. Fortunately, population size heterogeneity makes confinement more promising for both systems, increasing the required number of daily migration events for batches of 10 adults by 3-4 for translocations and by 1-2 for UD^MEL^.

## Discussion

The idea of using threshold-dependent gene drive systems to replace local populations of disease vectors with varieties that are unable to transmit diseases has been discussed for over half a century now, since Curtis (36) famously proposed the use of translocations to control diseases transmitted by mosquitoes. While Curtis had been primarily concerned with introducing and spreading genes into a population, as the technology nears implementation, confining them is also becoming a significant concern. As CRISPR-based homing gene drive technology edges closer to field application, concerns are increasingly being raised regarding the invasiveness of these systems (45, 46), and systems such as split drive (11), daisy drive (12) and threshold-dependent underdominant systems (7) are gaining interest, at least during the trial phase of population replacement technology (6). In this paper, we model the introduction of two drive systems, chromosomal translocations and UD^MEL^, that have been engineered in the laboratory and shown to display threshold-dependent spread (14, 15). While previous papers have described the population dynamics of these two systems in randomly-mating populations ignoring life history (14–16, 36), with one recent paper including spatial structure (18), we present the first analysis of these systems in a spatially-structured population including mosquito life history and reflecting a well-characterized landscape where field trials could conceivably be conducted (21).

Our results provide strong support for the use of translocations to implement confineable and reversible population replacement in structured *Ae. aegypti* populations. Regarding reversibility, translocations are preferable to UD^MEL^ as: i) they can be remediated through releases of non-disease-transmitting male *Ae. aegypti*, and ii) required releases sizes are achievable (~10 weekly releases at a coverage level of 50%). UD^MEL^ requires less effort to introduce into a population; but is much more difficult to remove once it has been introduced, requiring a very large number of both males and disease-transmitting females to be released. This highlights the benefit of a ~50% threshold for reversible population replacement: the symmetry allows both replacement and remediation to be achieved with similar effort. Extreme underdominance is another example of system with a 50% threshold (20, 47). Regarding confineability, translocations again outperform UD^MEL^ as ~16 daily migration events of batches of 5 *Ae. aegypti* adults are required for translocations to spread to the neighboring suburb of Trinity Park, while UD^MEL^ can spread to Trinity Park given only ~3 daily migration events (or 2-3 daily migration events for alternative model parameterizations). The true batch migration rate between suburbs is expected to be smaller than either of these (21), however the rate required for translocations to spread is highly unlikely to be reached, while the rate for UD^MEL^ is conceivable.

As with any modeling study, there are limitations inherent in our analysis. Several of the parameters we assumed to be constant here would indeed be dynamic in a real intervention scenario. At the genetic level, lab experiments suggest non-outbred individuals homozygous for the translocation had a fitness cost that largely disappeared once offspring were produced that were the product of at least one wild-type individual (14). Models fitted to data from UD^MEL^ drive experiments also suggested dynamic fitness costs that depended on the frequency of transgenic organisms in the population (15). Another recent modeling study highlights the possibility of toxin and antidote mutational breakdown for underdominance constructs; however this is expected to be impactful over a larger timescale than considered here (hundreds of generations) (48). At the ecological level, our model of *Ae. aegypti* life history (26), based on the lumped age-class model of Hancock & Godfray (27), assumes the existence of a constant equilibrium population size, and other constant ecological parameters, such as adult death rate and larval development times. These parameters have indeed been shown to vary in space and time, and in response to local mosquito density (49, 50), which our sensitivity analyses suggest could have significant impacts on release thresholds and gene drive outcomes (43) (Figure 5 & S4). At the landscape level, we have assumed a relatively homogenous distribution of mosquitoes per household, and movement rates between households that depend only on distance and household distribution. Extensive landscape heterogeneities have been shown to slow and alter the spread of *Wolbachia* (32, 51), and would likely impact the spread of translocations and UD^MEL^ as well. Future work that helps to characterize the environmental drivers of mosquito population dynamics will inform iterative model development to address this.

In conclusion, our analysis supports the use of translocations as a threshold-dependent drive system capable of spreading disease-refractory genes into structured *Ae. aegypti* populations in a confineable and reversible manner. If such a system were engineered in *Ae. aegypti*, it would be an excellent candidate for the introduction of disease-refractory genes during the trial phase of population replacement technology, or whenever a localized release were otherwise desired. As the technology nears implementation, further ecological work characterizing the density-dependencies, seasonality and spatial heterogeneities of *Ae. aegypti* populations will be essential to enhance model predictions in preparation for field trials.

## Materials and methods

We used the MGDrivE framework (26) (https://marshalllab.github.io/MGDrivE/) to simulate releases of adult *Ae. aegypti* males homozygous for one of two threshold-dependent gene drive systems - reciprocal chromosomal translocations or UD^MEL^ - in the community of Yorkeys Knob in Queensland, Australia. To simulate remediation, we modeled releases of wild-type adult *Ae. aegypti* into populations in Yorkeys Knob already fixed for the gene drive system. Houses receiving releases were randomly chosen from a uniform distribution. These were conserved within simulation runs, but varied between runs. To determine confineability, we simulated batch migration events from Yorkeys Knob (fixed for the gene drive system) to the neighboring community of Trinity Park (initially wild-type). The MGDrivE framework models the egg, larval, pupal and adult (male and female) mosquito life stages implementing a daily time step, overlapping generations and a mating structure in which adult males mate throughout their lifetime, while adult females mate once upon emergence, retaining the genetic material of the adult male with whom they mate for the duration of their adult lifespan. Density-independent mortality rates for the juvenile life stages are assumed to be identical and are chosen for consistency with the population growth rate in the absence of density-dependent mortality. Additional density-dependent mortality occurs at the larval stage, the form of which is taken from Deredec *et al.* (28). Full details of the modeling framework are available in the S1 Text of Sánchez *et al.* (26), and in the software documentation available at https://marshalllab.github.io/MGDrivE/docs/reference/. Parameters describing *Ae. aegypti* life history and the gene drive systems and landscape of interest are listed in Table S1.

The inheritance patterns for reciprocal chromosomal translocations (depicted in Figure 1A) and UD^MEL^ (depicted in Figure S1) are modeled within the inheritance module of the MGDrivE framework (26), and their impacts on female fecundity and adult lifespan are implemented in the life history module. The distribution of households in Yorkeys Knob, Trinity Park and Gordonvale were taken from OpenStreetMap (https://www.openstreetmap.org/) (Figure 1C). We implement the stochastic version of the MGDrivE framework to capture the randomness associated with events that occur in small populations, such as households, which serve as nodes in the landscape modeled here. In the stochastic implementation of the model, the number of eggs produced per day by females follows a Poisson distribution, the number of eggs having each genotype follows a multinomial distribution, all survival/death events follow a Bernoulli distribution, and female mate choice follows a multinomial distribution with probabilities given by the relative frequency of each adult male genotype in the population.

## Acknowledgements

The authors would like to thank Dr. Gregory Lanzaro, Dr. Yoosook Lee, Dr. Tomás León, and Ms. Partow Imani for discussions on *Aedes aegypti* life history and dispersal behavior. This work was supported by a DARPA Safe Genes Program Grant (HR0011-17-2-0047), awarded to OSA and JMM, and funds from the Innovative Genomics Institute, awarded to JMM.

## Author contributions

JMM and HMSC conceptualized the study. HMSC, JBB and SLW performed the experiments and visualized the results. GR and OSA contributed to the interpretation of the results. JMM wrote the first draft of the manuscript. All authors contributed to the writing of and approved the final manuscript.

---

**Figure S1.**
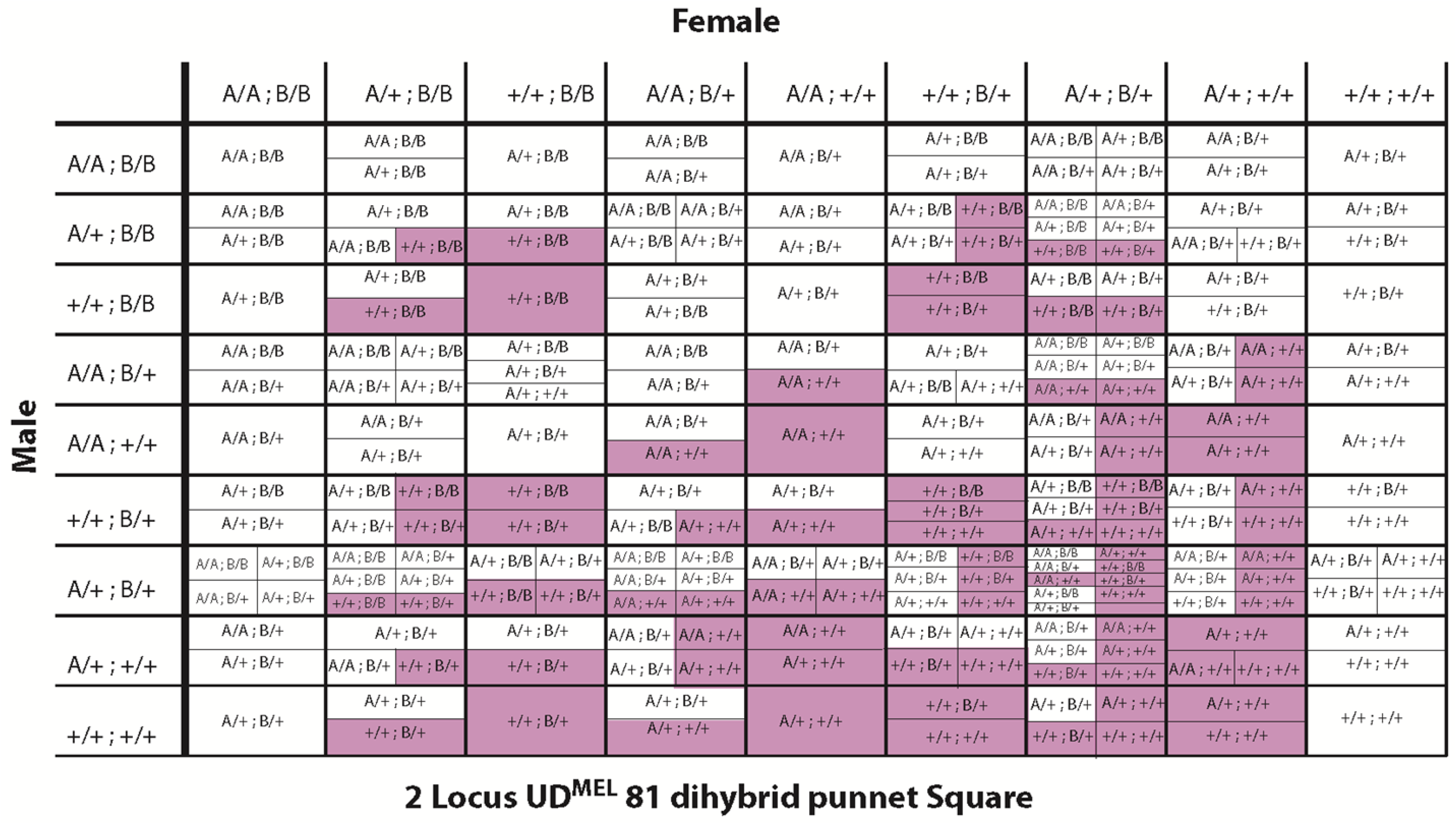
Complete inheritance pattern of UD^MEL^. UD^MEL^ is composed of two unlinked constructs (here referred to as A and B), each consisting of a maternally-expressed toxin and a zygotically-expressed antidote for the toxin on the opposite construct (see Figure 1B). The cross here represents matings between all nine possible parental genotypes (“+” represents the wild-type allele, and “A” and “B” represent alleles corresponding to the two UD^MEL^ constructs). Offspring lacking the antidotes to the maternal toxins produced by their mother are unviable (shaded). At high population frequencies, the selective advantage on the constructs, by virtue of the antidotes, outweighs the fitness load due to the toxins, and hence results in frequency-dependent spread.

**Figure S2.**
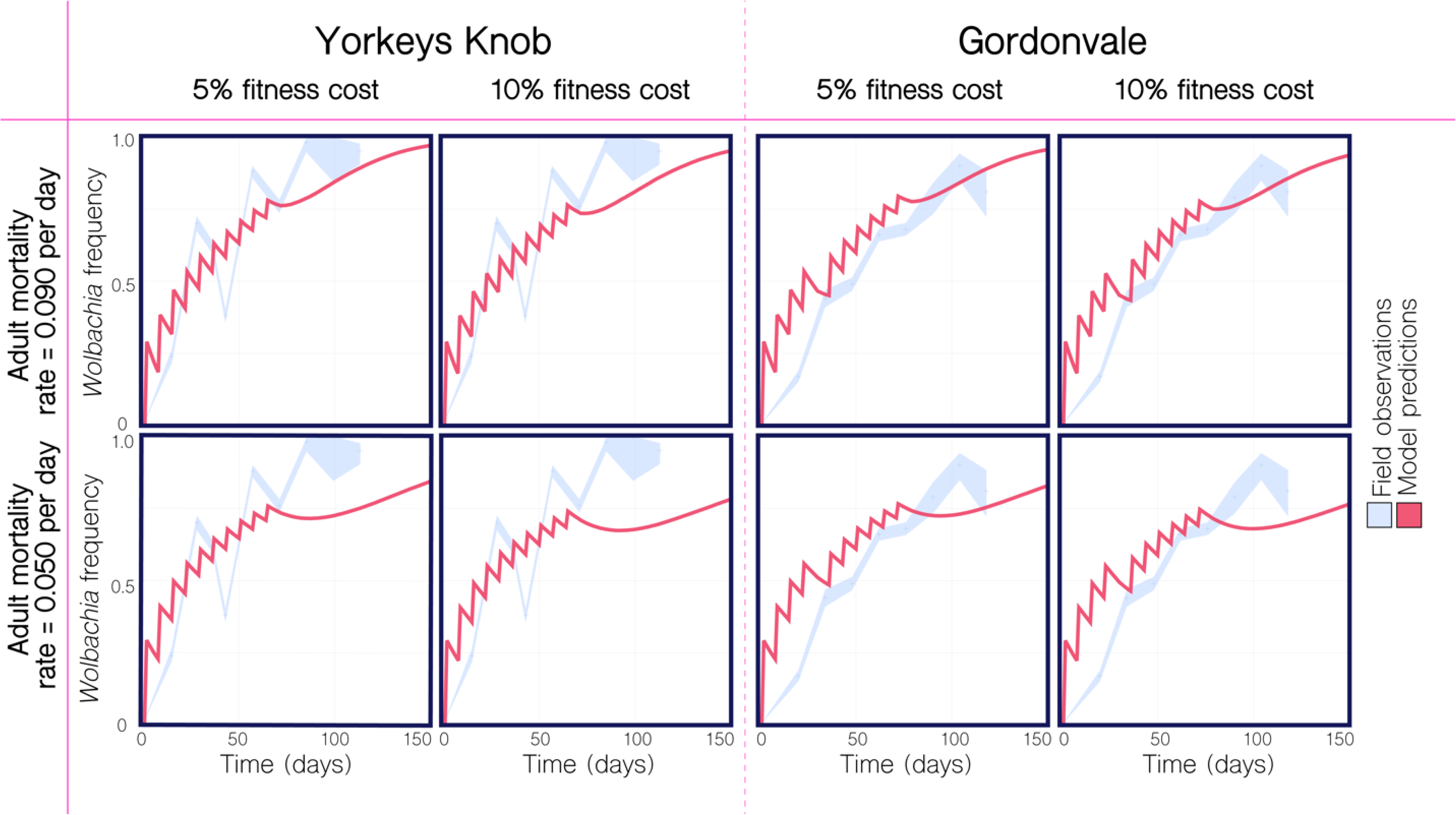
Comparison of model predictions to *Wolbachia* field trial data from Yorkeys Knob and Gordonvale, Australia for a variety of parameterizations. Field observations of *Wolbachia* population frequency are depicted in light blue, with 95% binomial confidence intervals based on the frequency and sample size reported for the 2011 field trials in Yorkeys Knob and Gordonvale. Model predictions are depicted for an analogous release scheme consisting of 20 *Wolbachia*-infected mosquitoes (10 female and 10 male) per household at a coverage of 30% over 10 weeks with the exception that, in Gordonvale, the fifth release was postponed by a week due to a tropical cyclone. Parameter values are listed in Table S1. Model predictions more closely match observed field data for a baseline adult mortality rate of 0.090 per day *(Top)*, as compared to one of 0.050 per day *(Bottom)*. Observations match predictions well for both 5% and 10% fitness costs associated with *Wolbachia*. Agreement between observations and predictions is strong, providing good validation for the modeling framework.

**Figure S3.**
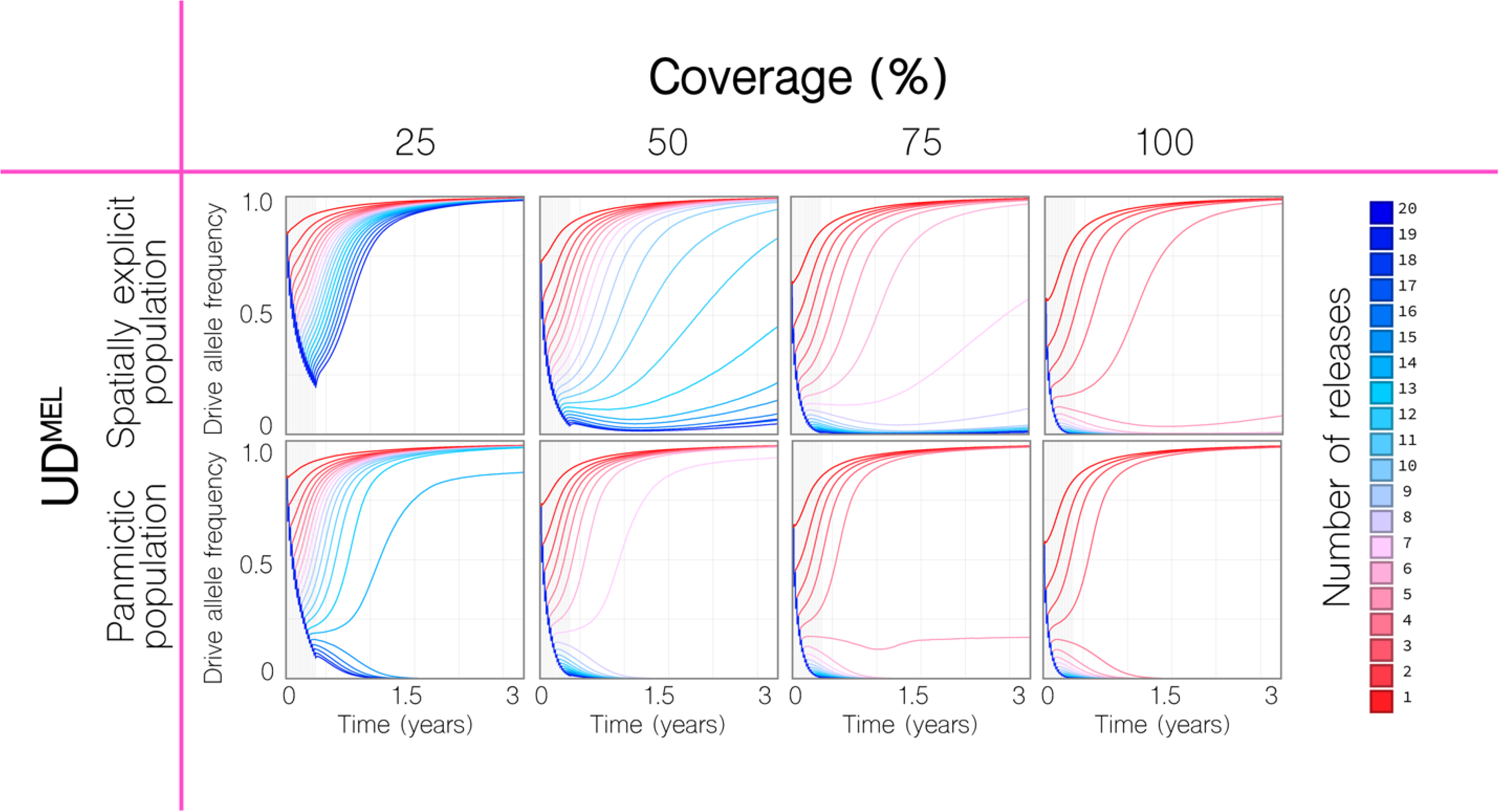
Remediation results for UD^MEL^ in spatially-explicit and panmictic populations. Time-series results are shown for a given number of weekly releases of 20 adult wild-type *Ae. aegypti* per household (10 female and 10 male) with the intent of remediation in the community of Yorkeys Knob (Figure 1C), and at given coverage levels, where coverage is the proportion of households that receive the releases. *(Top)* Remediation in the spatially-explicit population is extremely difficult. At a coverage level of 75%, nine weekly releases are required for a reduction in UD^MEL^ allele frequency over the first year; however, complete remediation is not possible even with 20 releases. Complete remediation is possible at a coverage level of 100% for 15 or more weekly releases. *(Bottom)* Remediation in the panmictic population is much less demanding. At a coverage level of 25%, it can be achieved with 16 or more releases, and at a coverage level of 50%, it can be achieved with nine or more releases.

**Figure S4.**
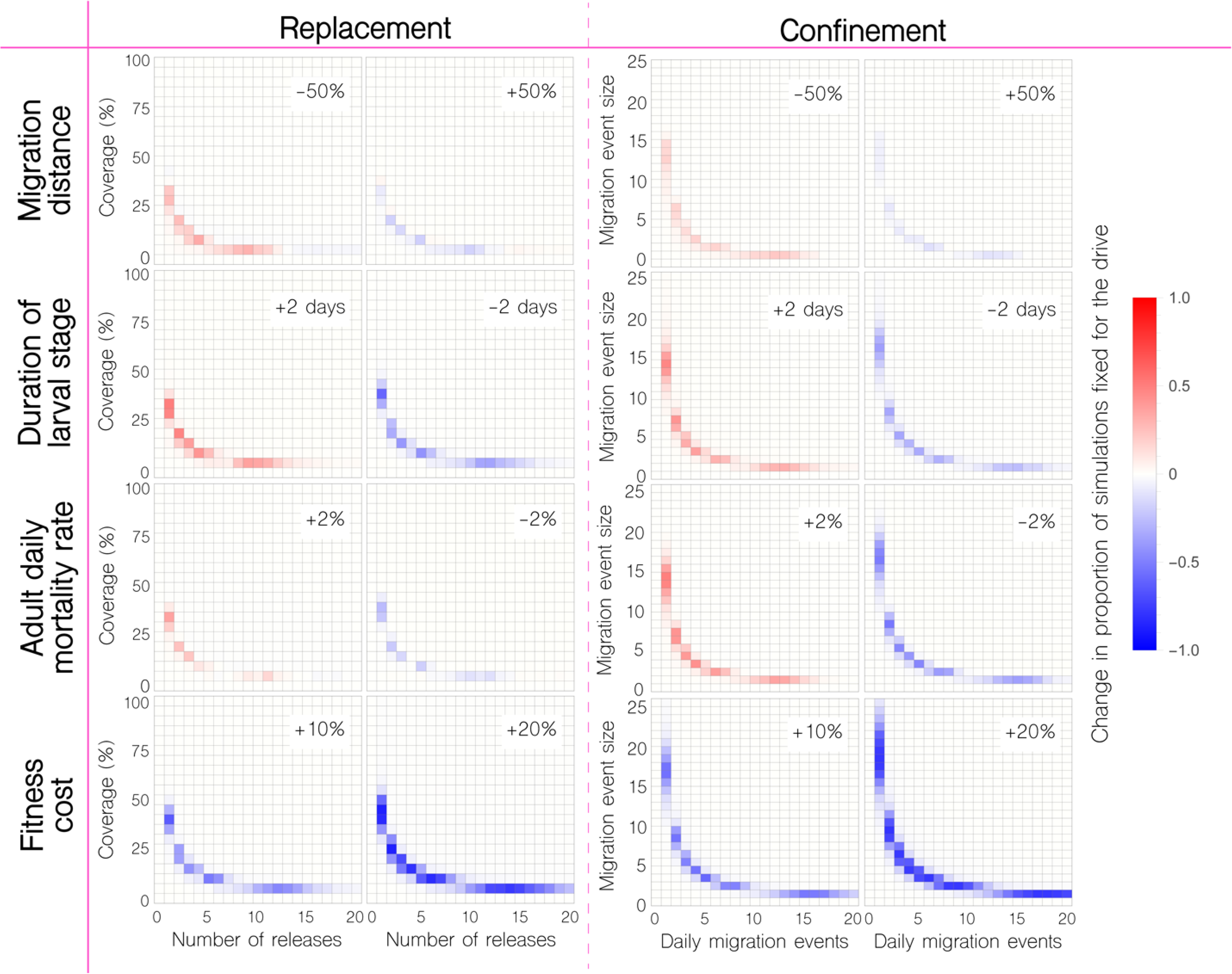
Sensitivity of model outcomes for replacement, remediation and confinement of UD^MEL^. Changes are depicted in the proportion of 50 stochastic simulations that result in fixation for replacement and confinement of UD^MEL^. Proportions are compared to those in the second row of Figure 4 as we vary: i) the mean dispersal distance of adult mosquitoes (+/− 50%), ii) the duration of the larval life stage (+/− 2 days), iii) the baseline adult mortality rate (+/− 2%), and iv) the fitness cost associated with being homozygous for the translocation (+10% or +20%). UD^MEL^ displays similar parameter sensitivities regarding fixation and batch migration outcomes as for translocations (Figure 5), with the exception that these outcomes are less sensitive to fitness costs, likely due to the fact that fitness is accommodated through a reduction in female fecundity rather than an increase in adult mortality.

**Figure S5.**
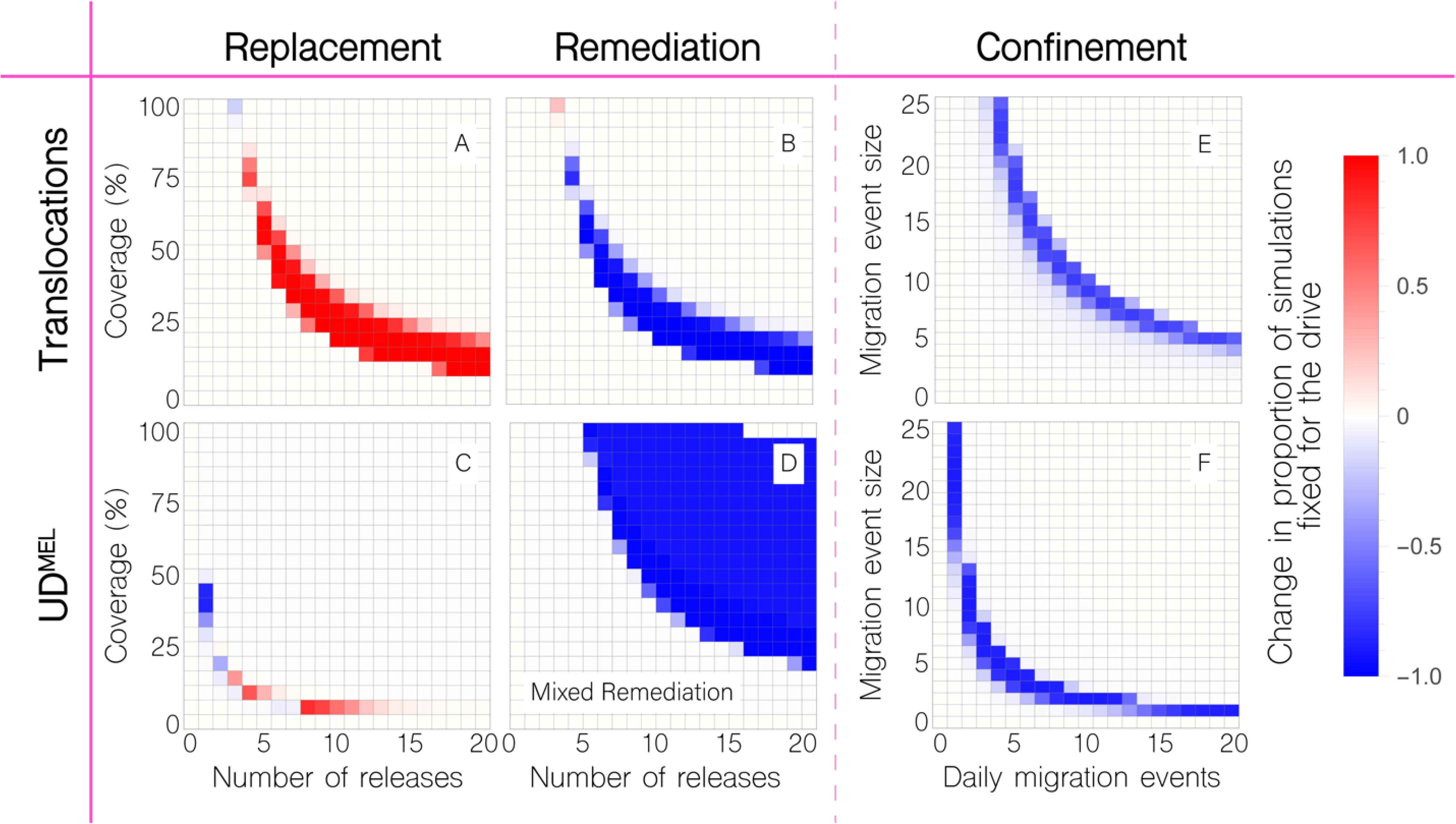
Sensitivity of model outcomes for translocations and UD^MEL^ comparing spatially-explicit and panmictic populations. Changes are depicted in the proportion of 50 stochastic simulations that result in fixation for replacement, remediation and confinement of translocations and UD^MEL^. Proportions are compared to those in Figure 4 as we simulate a model where Yorkeys Knob and Trinity Park are panmictic populations of equivalent size to their spatially-explicit versions. Introducing population structure greatly increases the release requirements to remediate UD^MEL^ (as seen in Figure S3), and substantially increases the release requirements for replacement or remediation of translocations. Invasion of a neighboring population, on the other hand, requires moderately fewer daily migration events in structured populations for both translocation and UD^MEL^.

**Figure S6.**
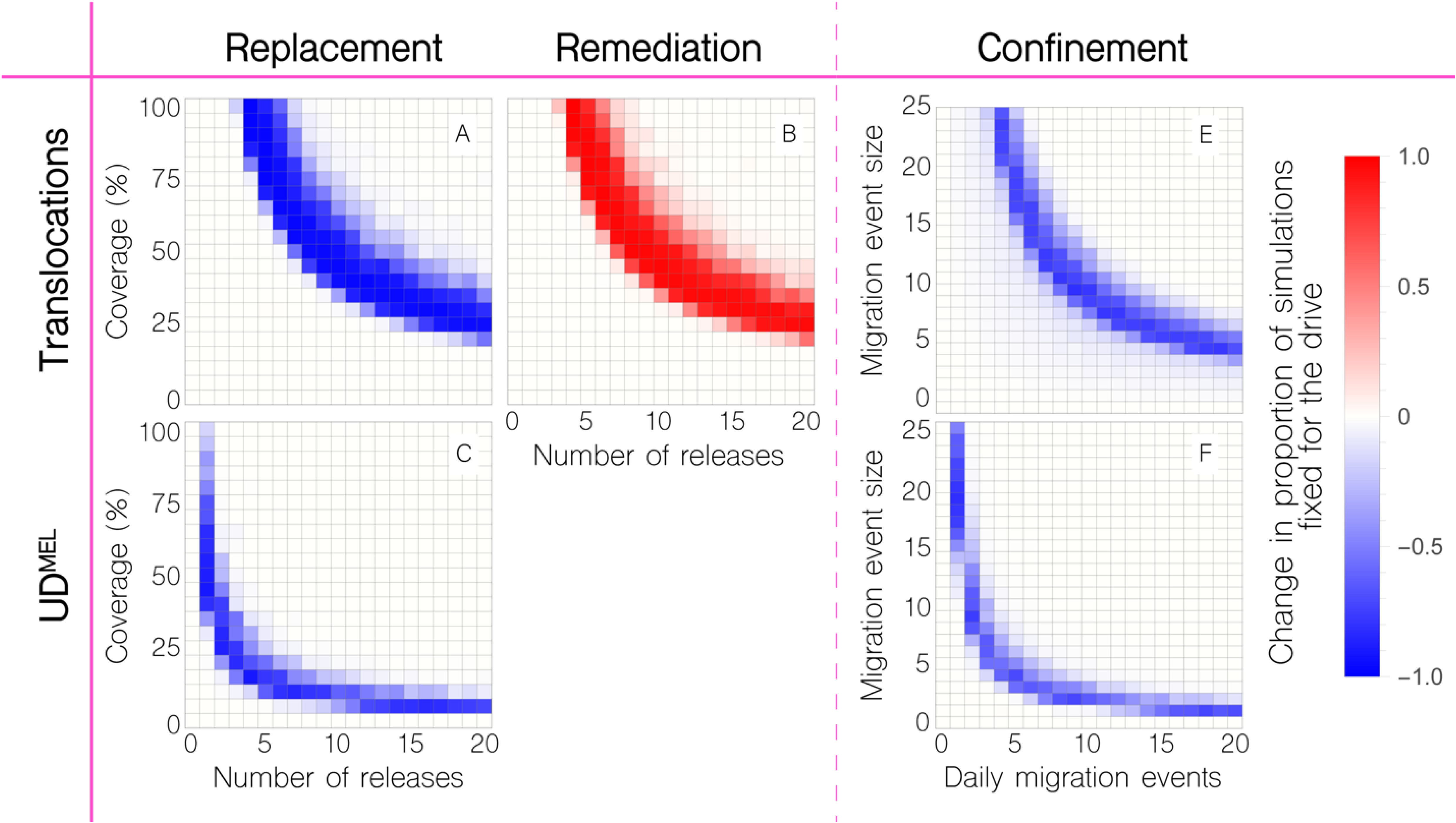
Sensitivity of model outcomes for translocations and UD^MEL^ comparing spatially-explicit populations with and without heterogeneity in household mosquito population size. Changes are depicted in the proportion of 50 stochastic simulations that result in fixation for replacement, remediation and confinement of translocations and UD^MEL^. Proportions are compared to those in Figure 4 as we simulate a model where household mosquito population size is distributed according to a zero-inflated, truncated exponential distribution with a mean of 15 adults, 55% of households having no mosquitoes, and none having more than 45 adults. Introducing household population size heterogeneity substantially increases release requirements for replacement and remediation with translocations, and marginally increases release requirements for replacement with UD^MEL^. Fortunately, population size heterogeneity makes confinement moderately more promising for both systems.

**Table S1.**
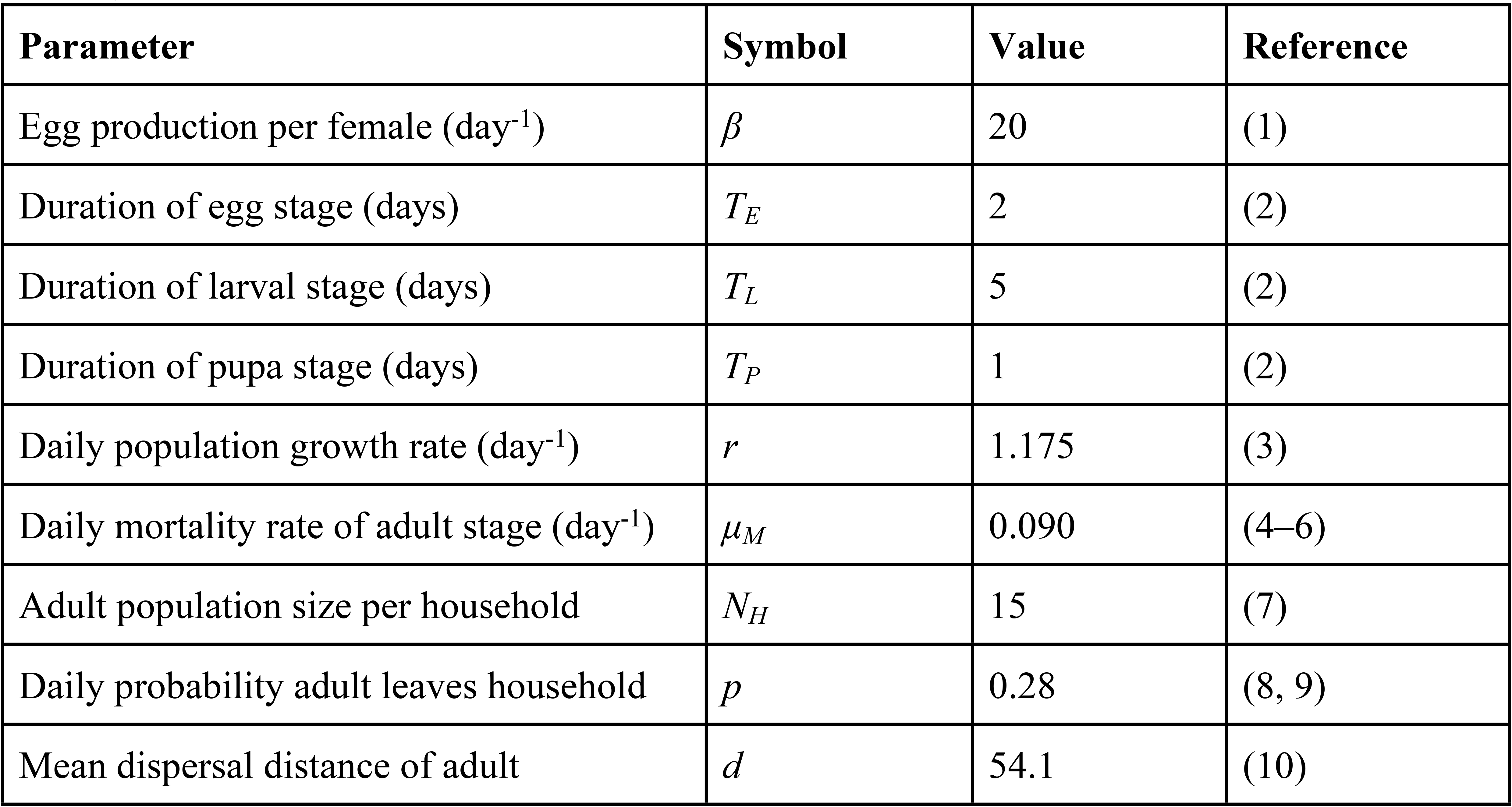
Life history, population size and movement parameters for *Aedes aegypti* in Cairns, Australia.

| Parameter | Symbol | Value | Reference |
| --- | --- | --- | --- |
| Egg production per female (day<sup>-1</sup>) | $\beta$ | 20 | (1) |
| Duration of egg stage (days) | $T_E$ | 2 | (2) |
| Duration of larval stage (days) | $T_L$ | 5 | (2) |
| Duration of pupa stage (days) | $T_P$ | 1 | (2) |
| Daily population growth rate (day<sup>-1</sup>) | $r$ | 1.175 | (3) |
| Daily mortality rate of adult stage (day<sup>-1</sup>) | $\mu_M$ | 0.090 | (4–6) |
| Adult population size per household | $N_H$ | 15 | (7) |
| Daily probability adult leaves household | $p$ | 0.28 | (8, 9) |
| Mean dispersal distance of adult | $d$ | 54.1 | (10) |

